# URB597 induces aggression in adult Lister Hooded rats

**DOI:** 10.1101/2022.03.09.483601

**Authors:** William G. Warren, Ed Hale, Eleni P. Papagianni, Helen J. Cassaday, Carl W. Stevenson, Christine Stubbendorff

## Abstract

The endocannabinoid system has been implicated in both social and cognitive processing. The endocannabinoid metabolism inhibitor, URB597, dose-dependently improves non-social memory in adult Wistar and Sprague Dawley rats, whereas its effect on social interaction (SI) is affected by both rat strain and drug dose. Lister Hooded rats consistently respond differently to drug treatment in general compared with albino strains. This study sought to investigate the effects of different doses of URB597 on social and non-social memory in Lister Hooded rats, as well as analysing the behavioural composition of the SI.

Males were tested for novel object recognition (NOR), social preference (between an object and an unfamiliar rat), social novelty recognition (for a familiar vs unfamiliar rat) and SI with an unfamiliar rat. URB597 (0.1 or 0.3 mg/kg) or vehicle was given 30 minutes before testing. During SI testing, total interaction time was assessed along with time spent on aggressive and explorative behaviours.

Lister Hooded rats displayed expected non-social and social memory and social preference, which was not affected by URB597. During SI, URB597 did not affect total interaction time. However, the high dose increased aggression, compared to vehicle, and decreased anogenital sniffing, compared to the low dose of URB597. In summary, URB597 did not affect NOR, social preference or social recognition memory but did have subtle behavioural effects during SI in Lister hooded rats.

These findings highlight the importance of considering strain as well as the composition of behaviour when investigating drug effects on social behaviour.

## Introduction

Adaptive social interaction requires the correct interpretation of social cues and subsequent adjustment of behaviour to fit the situation. Both social processing and management of social behaviour are impaired in psychiatric disorders, strongly impacting the ability to function in a social context (Grady and Keightley 2002, Weightman et al. 2014, Green et al. 2015). Given the prevalence of social impairment in mental illness, it is important to understand how current and novel treatments affect social cognition and behaviour across patient populations. Current antipsychotic and antidepressant drugs are not always effective and can induce side effects in a considerable number of patients, even after showing efficacy in preclinical trials (Tawa and Murphy 2013, Singewald et al. 2015). This is likely due to the lower genetic variability in laboratory rodents, compared to humans. The disparity in drug efficacy between preclinical and clinical studies demonstrates the importance of modelling a more diverse population in preclinical studies. Pharmacological treatments are commonly tested in albino rat strains (Wistar or Sprague-Dawley). However, drugs often produce different behavioural effects when tested on pigmented strains such as Long Evans and Lister hooded (LH) rats (Horowitz et al. 1999, Johnson and Mitchell 2003, McDermott and Kelly 2008, Hong et al. 2011, Neeley et al. 2011). This is possibly caused by neurophysiological differences between rat strains (Johnson and Mitchell 2003, Hong et al. 2011, Neeley et al. 2011). Considering strain when designing preclinical studies may provide an opportunity to model variation in drug response in the human population.

The endocannabinoid (eCB) system has received increased attention as a potential target for treating a range of psychiatric disorders (Batalla et al. 2019, Papagianni and Stevenson 2019, Warren et al. 2021, Zou et al. 2021). The eCB system is comprised of two receptors (cannabinoid receptor type 1 & 2 [CB1R & CB2R]) and their endogenous ligands (anandamide [AEA] and 2-arachidonoylglycerol [2-AG]), along with their respective metabolic enzymes (fatty acid amide hydrolase [FAAH] and monoacylglycerol lipase [MAGL]) (Blankman and Cravatt 2013). CB1R is preferentially expressed in the central nervous system, whilst CB2R is found more abundantly in the periphery. 2-AG has a higher affinity for CB1R and CB2R than AEA, whilst AEA shows greater affinity for transient receptor potential vanilloid 1 (TRPV1;(Ligresti et al. 2016). One mode of eCB transmission is via retrograde signalling, where 2-AG mediated activation of pre-synaptic CB1Rs leads to the inhibition of neurotransmitter release (Kano et al. 2009). Alternatively, post-synaptic activation of TRPV1 and CB1R can increase downstream neural activity; this is largely driven by AEA. The localisation of FAAH and MAGL correspond with the putative signalling mechanisms of 2-AG and AEA. FAAH inhibits post-synaptic signalling by metabolising AEA into arachidonic acid and ethanolamine, whilst MAGL metabolises 2-AG into arachidonic acid and glycerol in the pre-synaptic terminal (Cravatt et al. 1996, Dinh et al. 2002). The FAAH inhibitor URB597 selectively inhibits the metabolism of the AEA, thereby increasing its tone and availability to act on CB1R (Fegley et al. 2005).

The eCB system modulates social behaviour, cognition and memory (Terranova et al. 1996, Marsicano et al. 2002, Varvel et al. 2005). URB597 improves non-social memory in albino rat strains (Hasanein and Teimuri Far 2015, Hlavacova et al. 2015), whereas lower doses of URB597 increase and higher doses decrease social interaction in these strains (Manduca et al. 2015, Matricon et al. 2016). The evidence for these opposing behavioural outcomes from different URB597 doses suggests that social behaviour may be particularly sensitive to manipulation by CB1R signalling. However, the effects of URB597 on memory and social behaviour have not been tested in pigmented rat strains, such as LH. When examining drug effects on social behaviour, it is important to consider changes not only to the sum but also to the quality of social interaction. While the CB1R agonist WIN 55,212-2 reduced overall social interaction in Wistar rats, this reduction was driven by a decrease in “following” and “anogenital sniffing” of the conspecific (Schneider et al. 2008). Drug-induced changes to the expression of aggression is particularly interesting when examining the impact of eCBs on social behaviour. Violent aggression (self-directed or towards others) can be a major obstacle for treatment of psychiatric patients (Pekurinen et al. 2017, Reknes et al. 2017, Pitts and Schaller 2021). Most studies examining the effects of eCBs on aggression have found cannabinoids to ameliorate expressed aggression, but some reported increased aggression in response to elevated brain AEA, with a possible link between individual differences in the level of trait aggression and drug effects (Sulcova et al. 1998, Kolla and Mishra 2018, Chang et al. 2021). However, the effects of URB597 on overall distribution of social behavioural components and aggression remain to be elucidated. Here we characterise, for the first time, the effect of URB597 on non-social and social novelty recognition as well as its effect on social interaction and aggression in male LH rats.

## Methods

### Animals

The study used 48 male Lister Hooded rats (Charles River UK), weighing between 250g-300g upon arrival. Individually ventilated cages (maintained at a constant temperature of 23 ± 0.5°C) each housed 4 rats, with a light/dark cycle of 12h:12h (lights on at 7am). Food and water were available *ad libitum* whilst husbandry adhered to the principles of laboratory animal care. Each animal was tested at approximately the same time each day. Procedures used in the experiments had ethical approval from the institution’s ethics committee and adhered to the UK Animals (Scientific Procedures) Act 1986 (Home Office Project Licence 30/3230). The animals were tested as two temporally separate cohorts of 24 animals (n=8 for each treatment group in each cohort). The timeline and details of the tests in each cohort is depicted in Figure 1A.

**Figure 1.**
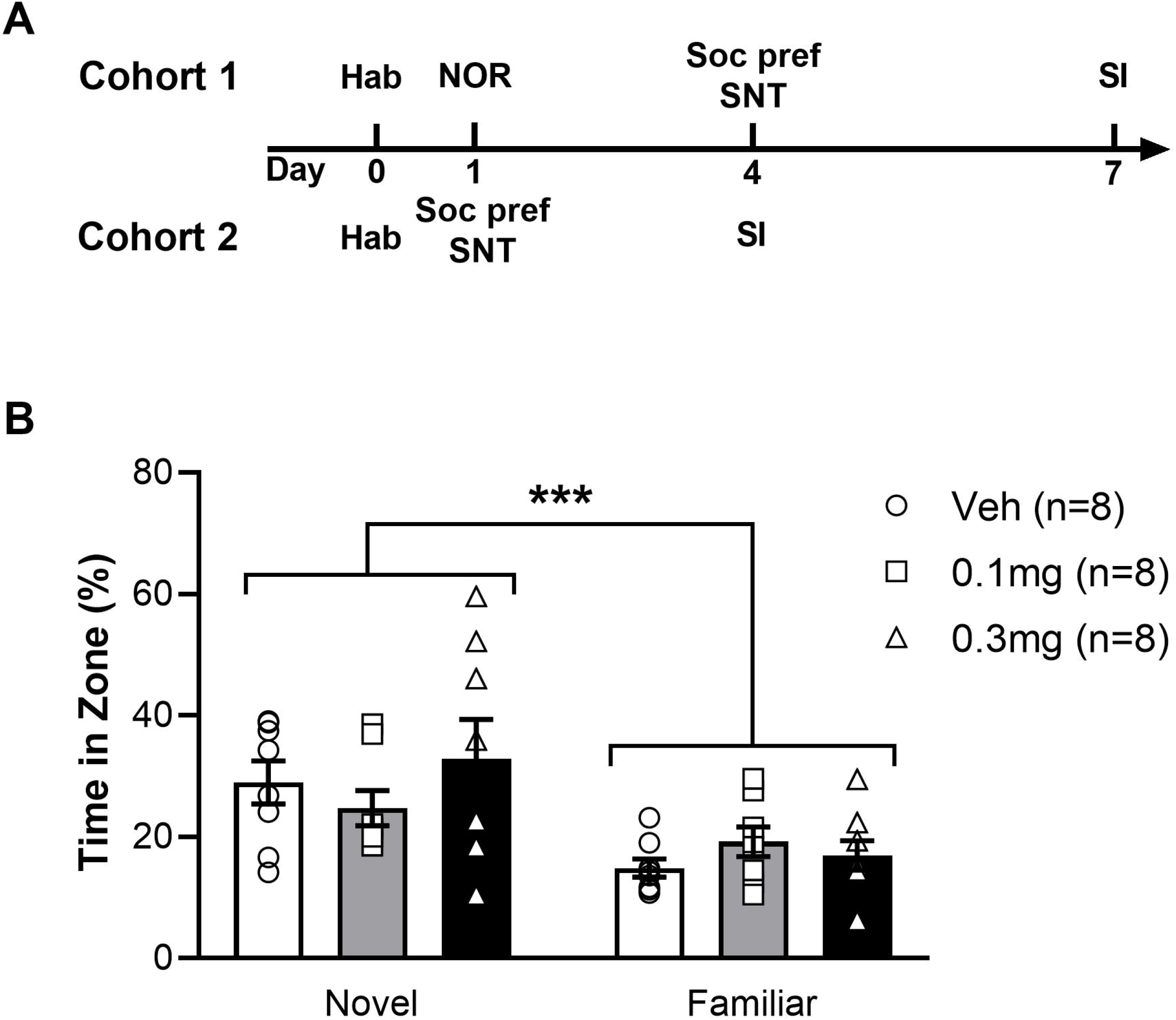
A. Timeline of experiments in cohort 1 and 2. B. Average percentage time (+/− SEM) spent exploring the novel or familiar object zone in LH rats treated with vehicle (white bar), 0.1 mg/kg URB597 (grey bar) and 0.3 mg/kg URB597 (black bar), with individual values displayed as circles, squares and triangles, respectively. All treatment groups preferred the novel to the familiar object, while NOR was not affected by URB597. Error bars represent +/− SEM. *** *p* = 0.0002

### Drugs

URB597 (0.1 and 0.3 mg/kg; Sigma-Aldrich, UK) was dissolved in 5% polyethylene glycol (Fluka Chemicals, Switzerland), 5% Tween 80 (Sigma-Aldrich) and 90% saline; doses and protocols were based on previous findings (Hasanein and Teimuri Far 2015, Hlavacova et al. 2015, Manduca et al. 2015, Matricon et al. 2016). Rats received either one of the URB597 doses or vehicle per experiment. All injections were administered intraperitoneally (1 ml/kg) and were given 30 minutes before testing.

### Apparatus and Materials

All experiments took place in an open field arena (L: 100cm, W: 100cm) above which a camera was positioned to record activity. Wire mesh cages were used to contain conspecifics in the social novelty test (SNT). All videos were recorded using EthoVision software (Noldus, the Netherlands).

### Habituation

Day 0 – Rats were habituated to the open-field for 10 minutes. The time spent in the corners of the open field was recorded and the rats’ preference for the corners that would contain stimuli in the following tests (see below) was calculated. This information was later used to counterbalance the placement of novel objects/conspecifics in the novel object recognition (NOR) and SNT experiments.

### Novel Object Recognition

Day 1 – The first phase was prepared by placing two identical objects (tin cans) in opposite corners of the open field. Test rats were individually placed in a corner of the open field, equidistant from the tin cans and allowed to explore both objects. After 10 minutes the rats were returned to a holding cage for 2 minutes while one of the tin cans was replaced with a glass jar (novel object), and an identical tin can replaced the one previously used (familiar object). In the second phase, the test rats were re-introduced to the arena for another 10 minutes before being returned to their home cage. Objects and the test arena were cleaned with 70% ethanol between trials. EthoVision was used to assess time spent within the two 25×25cm corner zones containing an object.

### Social Novelty Test

Day 1 or 4 – The SNT was adapted from Seillier and Giuffrida (2016) and composed of two separate consecutive phases, depicted in Figure 2A and 2B. The first phase was a social preference test, where two wire mesh cages were placed in opposite corners of the arena. One cage was empty, whilst the other held an unfamiliar, weight-matched male conspecific. Test rats were placed in the corner of the arena, equidistant from both cages. The test rat then explored the open field for 10 minutes, after which they were removed to a holding cage. The time spent in the vicinity of the cages was recorded with EthoVision software (Noldus, the Netherlands).

**Figure 2.**
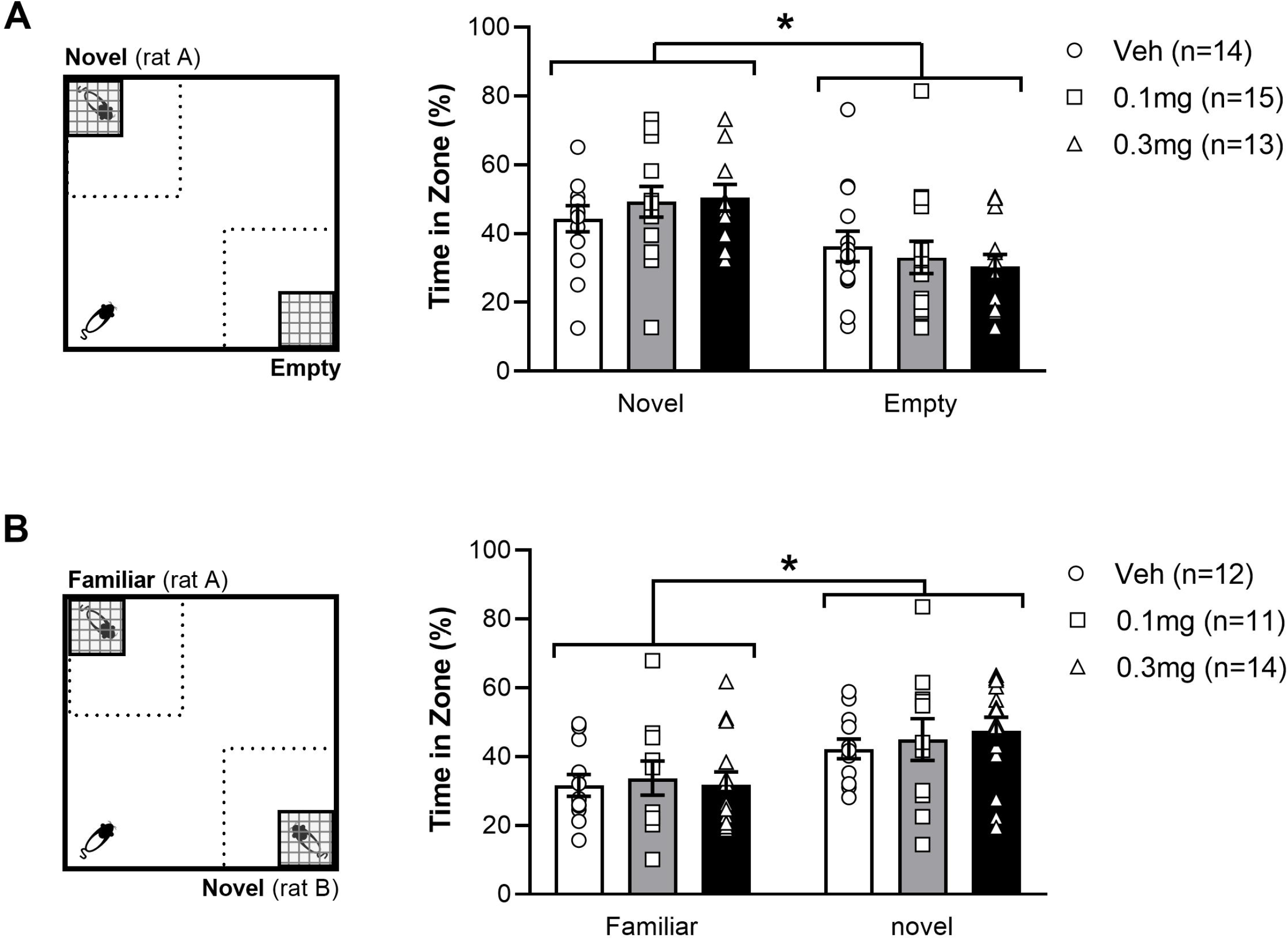
Left: Experimental set up of Social Preference test (A) and Social Novelty Test (SNT) (B). The dotted square represents the 40×40cm interaction zones. Right: Average percentage time (+/− SEM) spent exploring the stimulus zones during the social preference test (A) and SNT (B) in LH rats treated with vehicle (white bar), 0.1 mg/kg URB597 (grey bar) and 0.3 mg/kg URB597 (black bar), with individual values displayed as circles, squares and triangles, respectively. A. All treatment groups preferred the social stimulus (novel rat) to the non-social stimulus (empty cage), while social preference was not affected by URB597 treatment. B. All treatment groups spent more time exploring a novel unfamiliar, compared with a familiar conspecific, while social novelty preference was not affected by URB597 treatment. Error bars represent +/− SEM. ** *p* < 0.04

The second phase of the experiment tested the rat’s social memory in a manner akin to the NOR. Immediately after the social preference test, a second unfamiliar weight-matched male conspecific i.e. (novel rat) was placed in the previously empty cage and then the cage was returned to its previous position. The stimulus rat used in the social preference test phase remained in its mesh cage in the same position and was now considered a familiar rat. Test rats then explored for another 10 minute period before being placed back in their home cage. EthoVision calculated the time that test rats spent in the 40×40cm interaction zone in the corners containing the cages.

### Social Interaction

Day 4 or 7 – Two weight-matched unfamiliar rats received the same drug treatment and were simultaneously placed in opposite corners of an open-field for 10 minutes. The rats were then free to interact during this time and behaviour was scored manually by an observer who was blind to the animals’ treatment. Total SI was defined as time engaged in any of the following behaviours: sniffing conspecific; following; crawling over/under; and aggression. To select behavioural components for further analysis, all social behaviours were identified and scored in a subset of videos initially using behavioural categories found in the literature (Watson et al. 2016, Kohli et al. 2019). Based on this initial exploratory analysis, head/body sniffing, anogenital sniffing and aggression (boxing, pinning or biting) were selected and scored in all pairs to test for drug effects. The social behaviour of each rat was scored separately and the average between interacting pairs then taken as the data output from the trial.

### Data Analysis

In the NOR and SNT, time spent in each zone over the course of the sessions was converted to a percentage. In the NOR, data from the full 10 minute session were used. However, only data from the first 5 minutes of the SNT were used because many of the rats moved the stimulus cages from their original corners later on during testing. Rats that moved the cages prior to the 5-minute mark were removed from the analysis. Differences between groups and the zones that the rats explored were statistically assessed with a two-way analysis of variance (ANOVA). Drug treatment formed the between-subject factor, whilst zone formed the within-subject factor. For the SI behavioural data, one vehicle treated rat and one rat treated with 0.3mg/kg URB597 were removed as statistical outliers based on Grubbs test (α=0.05) and all data from these rats was omitted from the analysis. Differences between groups were assessed using one-way ANOVA, with treatment as the between-subject factor.

Multiple comparisons tests were conducted using Tukey’s multiple comparisons test. All data are expressed as mean ± SEM. P values <0.05 were deemed significant. All videos and data analysis are available at https://osf.io/zsm8f/

## Results

### URB597 did not affect memory in the novel object recognition test

To assess the impact of URB597 on non-social cognition, we tested our LH rats in the NOR paradigm (Figure 1). All treatment groups spent a greater amount of time investigating the novel object zone than the familiar object zone (F (1, 42) = 16.66; *p* = 0.0002), demonstrating non-social novelty recognition and memory in LH rats, which is in accord with previous findings (Cyrenne and Brown 2011, Renard et al. 2013). NOR was not affected by URB597 in our LH rats, as indicated by the lack of significant main effect of treatment (F (2, 42) = 0.4736; *p* = 0.6260) or zone x treatment interaction (F (2, 42) = 1.226; *p* = 0.3037) in our LH rats.

### URB597 did not affect social preference or social memory in a social novelty task

The effect of URB597 on preference for social over non-social stimuli was tested by allowing the rats to explore an empty cage (non-social stimulus) and a cage containing an unfamiliar conspecific (social stimulus) in opposite corners of an open field (Figure 2A). All treatment groups spent a greater amount of time in the zone with social stimuli than in the zone with the non-social stimuli (F (1, 39) = 9.553; *p* = .0037) However, we found no effect of URB597, as indicated by the lack of significant main effect of treatment (F (2, 39) = 0.3668; *p* = 0.6953) or a zone x treatment interaction (F (2, 39) = 0.5362; *p* = 0.5892), suggesting that URB597 does not alter social preference in LH rats. We then used the SNT to examine if the lack of drug effect in LH rats observed in the NOR test was specific to non-social memory (Figure 2B). Similar to the observation in the NOR test, all treatment groups spent a greater amount of time in the zone with the novel conspecific than the (now familiar) conspecific used in the social preference phase (F (1, 34) = 7.235; *p* = 0.0110) but social novelty preference was not affected by URB597, as indicated by the lack of significant main effect of treatment (F (2, 34) = 1.636; *p* = 0.2096) or zone x treatment interaction (F (2, 34) = 0.1195; *p* = 0.8878). These results indicate that novelty preference was consistently unaffected by URB597 in LH rats, regardless of the social or non-social nature of the novel stimuli.

### URB597 did not affect total social interaction but increased aggressive behaviours

URB597 has previously been shown to have both dose- and strain-specific effects on total social interaction in albino rat strains (Manduca et al. 2015, Matricon et al. 2016), therefore we decided to investigate the effects of URB597 on social interaction in LH rats (Figure 3A). Unlike the reported effects in albino strains, we found no effect of either URB597 dose on total SI in LH rats, as indicated by the lack of significant main effect of treatment (F (2, 19) = 0.8445; *p* = 0.4453). This suggests that, overall, URB597 has less effect on SI in LH rats than albino strains. However, we noticed that specific social behaviour components, namely head/body sniffing, anogenital sniffing and aggressive behaviour (boxing, pinning and biting), were more expressed in some pairs than others. We therefore assessed the effect of URB597 on these behaviours (Figure 3B). Whereas URB597 treatment did not alter the amount of head/body sniffing (F (2, 19) = 1.010; *p* = 0.3829), LH rats treated with 0.1 mg/kg of URB597 spent more time engaged in anogenital sniffing, compared with 0.3 mg/kg of URB597 (F (2, 19) = 3.754; *p* = .0423; post hoc test *p* < 0.05). Finally, the higher dose of URB597 increased the amount of aggression, compared to vehicle treatment (F (2, 19) = 5.336; *p* = 0.0145; post hoc test *p* < .05). Taken together, these findings show that while URB597 did not alter total SI in LH rats, this drug shifted SI towards more dominance- and aggression-related behaviours.

**Figure 3.**
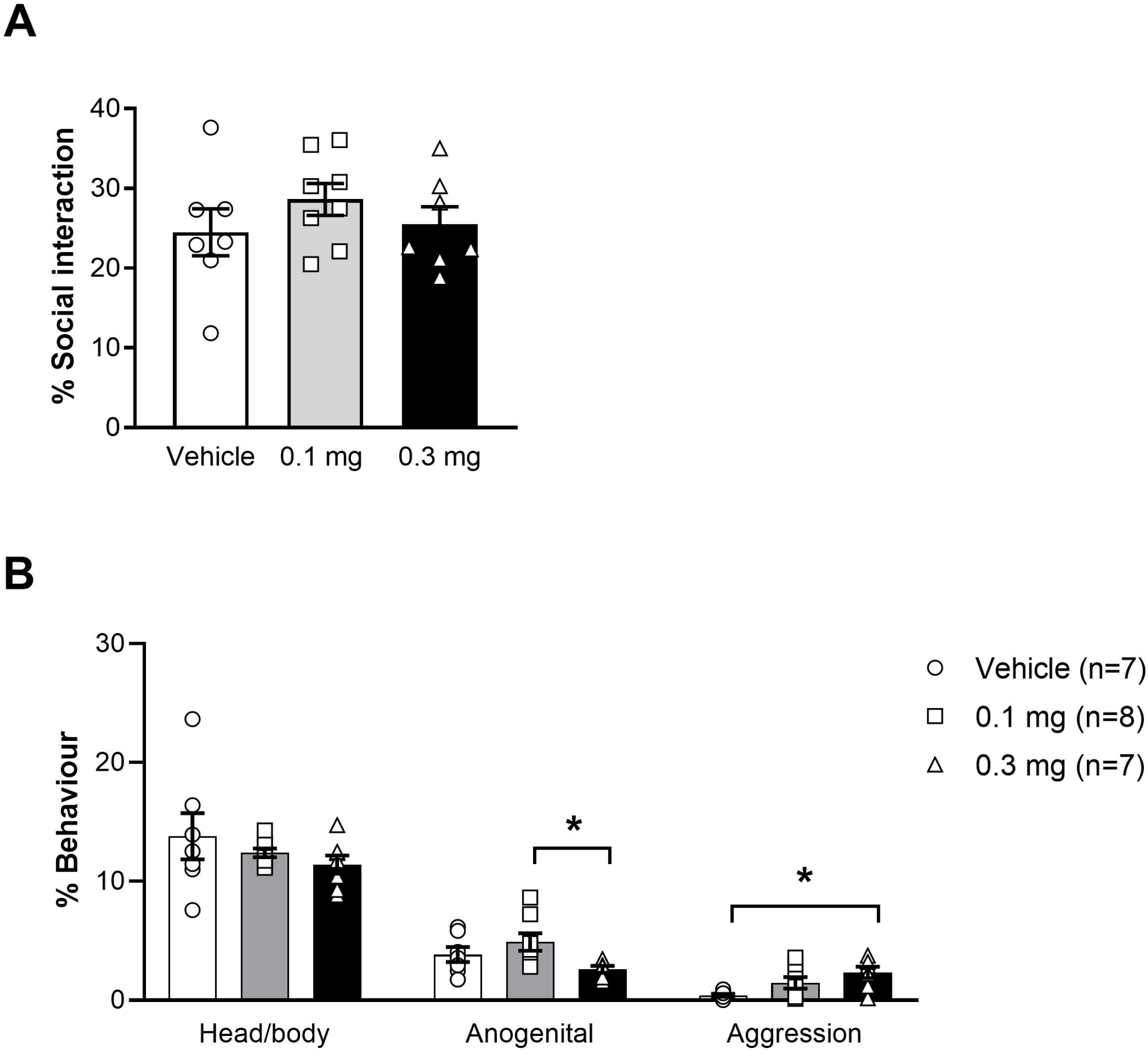
Average percentage time (+/− SEM) spent engaged in social interaction (A) or specific social behaviours (B) in LH rats treated with vehicle (white bar), 0.1 mg/kg URB597 (grey bar) and 0.3 mg/kg URB597 (black bar), with individual values displayed as circles, squares and triangles, respectively. A. URB597 had no effect on total SI in LH rats. B. Rats treated with 0.3 mg/kg URB597 spent less time engaged in anogenital sniffing, compared with rats treated with 0.1 mg/kg URB597 (middle), and more time engaged in aggressive behaviours, compared with vehicle treated rats (right). Error bars represent +/− SEM. * *p* < 0.05

## Discussion

### URB597 does not affect recognition memory in LH rats

While the effects of URB597 on non-social cognition and social behaviour have previously been assessed in albino rat strains. We tested its efficacy in LH rats and found that this strain responded differently compared with albino strains. Previous work found 0.3mg/kg URB597 to improve NOR in male Wistar as well as Sprague-Dawley rats (Hasanein and Teimuri Far 2015, Hlavacova et al. 2015), whereas NOR performance in male LH rats in the present study was not affected by either dose of URB597. Chronic treatment with the synthetic cannabinoid CP55,940, impairs NOR in Wistar but not in LH rats (Renard et al. 2013), suggesting that LH rats may be less sensitive to manipulation of cannabinoid receptor signalling, compared with albino rat strains.

In the current study, LH rats showed a preference for social versus non-social stimuli and also displayed social novelty recognition. Previous work, using a 10 minute three-chamber paradigm, have reported social preference but lacking social novelty recognition in LH males (McKibben et al. 2014). We focussed on the first 5 minutes of the exploration, when the stimuli are most novel, which may explain why we were able to detect social novelty recognition. We found not effect of URB597 on neither social preference nor social novelty recognition. While the effect of URB597 on social preference, to our knowledge, has not been examined in albino rat strains, CP55,940 dose-dependently suppressed social preference in male Wistar rats (Seillier and Giuffrida 2016), and cannabidiol decreased social recognition memory (Deiana et al. 2015), further suggesting increased sensitivity to cannabinoid modulation in albino strains, compared with LH rats. The SNT is comparable to the NOR in its setup and the combined use of these two tests enables the comparison of social and non-social components of novelty seeking and memory processing in rodents. Future work examining the effect of URB597 on social preference and in the social memory in albino strains may reveal whether the strain-dependent differences observed in NOR also manifest when the familiar and unfamiliar objects are replaced with conspecifics. LH rats display higher levels of baseline locomotor activity and have superior vision compared with albino strains, which may contribute to the difference in drug-induced behavioural output compared to albino strains (McDermott and Kelly 2008, Ihalainen et al. 2016). However, WIN 55,212-2, did not affect locomotion in neither male SD nor pigmented (LH or Long Evans) rats (Deiana et al. 2007), suggesting that locomotor effects alone cannot explain strain-differences in response to cannabinoid treatment. In addition, in an experiment with intravenous self-administration, both LH and Long Evans rats acquired stable WIN 55,212-2 self-administration behaviour whereas SD rats did not, further suggesting inherent differences between albino and pigmented rat strains in the response to cannabinoids (Deiana et al. 2007).

### URB597 increases aggression in LH rats

We found no effect of URB597 on the total sum of social interaction in LH rats, which is in contrast to findings in albino strains. 0.1mg/kg URB597 increased total SI in Wistar but not in Sprague-Dawley rats (Manduca et al. 2015), whereas the higher dose of 0.3mg/kg decreased SI in Wistar rats (Matricon et al. 2016). These findings further support an effect of strain in the response to URB597 on social behaviour.

We also quantified the sum of different behavioural components expressed during the SI test and found more anogenital sniffing in pairs treated with 0.1mg compared with 0.3mg of URB597. As the drug-induced shift in anogenital sniffing was not significantly different from behaviour in the vehicle-treated rats, interpretation of this result can only be speculative. Previous work reports that subordinate Long Evans rats decrease the amount of anogenital sniffing after direct confrontation by a dominant rat (Wesson 2013) and a link between anogenital sniffing behaviour and position within the dominance hierarchy has also been observed in mice (Lee et al. 2019). It is possible that the shift in anogenital sniffing in our LH rats is associated with assertion of dominance between the unfamiliar male rats. As our LH rat pairs were not re-tested to allow a dominance hierarchy to be established and assessed, it is beyond the scope of this study to draw conclusions on the role of URB597 on dominance in LH rats. However, our finding suggests that drug-induced changes to dominance-related behaviour is relevant to consider when testing the effects of eCB-modulating drugs

We observed a greater amount of time spent engaging in aggressive behaviour in pairs treated with 0.3mg URB597 compared with vehicle. To our knowledge, the effect of URB597 on aggression has not been assessed in rats before. However, in high aggressive mice, URB597 infusion into ventral hippocampus reduced aggressive behaviour in the SI (Chang et al. 2021) whereas male Syrian hamsters showed no change in defensive aggression in response to URB597 when tested as intruders in the resident/intruder paradigm (Moise et al. 2008). In addition, a study in male intruder mice in the resident/intruder paradigm reported that AEA increased territorial aggression in low-aggressive mice, while lowering territorial aggression in high-aggressive mice (Sulcova et al. 1998). Taken together, these findings suggest that pharmacological modulation of AEA alters the expression of aggressive behaviour across species, but that aggression may also be influenced by the testing parameters as well as individual differences in trait aggression. When interpreting observations of increased aggression, it is important to consider that aggression in itself is not maladaptive and plays a crucial role in survival. Neither is aggression necessarily violent (e.g. in the context of competition) but when aggression manifests out of context and/or out of proportion to the situation it can be considered maladaptive (Miczek et al. 2007, de Boer 2018). In our SI paradigm, the rats entered the arena together and, therefore, the observed increase in aggression cannot be defined as either resident-like territorial or intruder-like defensive aggression, nor can the increased aggression be defined as maladaptive (excessive) or adaptive, e.g. competitive aggression. The observed increase in aggression manifested specifically in more boxing, pinning and biting with no injuries (puncture of skin) to either rat. While such aggressive display can be considered mild, our findings do not provide detail on the adaptive or maladaptive nature of increased aggression induced by URB597 in LH rats. Like most laboratory rodent strains, LH rats are generally low-aggressive, and therefore, our findings do not provide information about how URB597 affects aggression in a high-aggressive rat strain. However, our findings demonstrate that changes in eCB availability can alter levels of aggression. Furthermore, our observations demonstrate the importance of investigating drug-related changes to social behaviour in more detail than merely the total sum of behaviour.

### Implications of strain-dependent differences: Nuisance or opportunity?

One possible conclusion from the data on behavioural differences between albino and pigmented stains is that when designing experiments, it is important to choose a strain appropriate for the planned testing regime. However, another direction would be to embrace strain-dependent differences in the design and interpretation of experiments to model population heterogeneity, which could increase the translational value of such research. Investigating strain differences may reveal neurobiological differences between treatment receptive and non-receptive individuals that are relevant to understanding variability in treatment response in patients. Albino rat strains show inherent variations in brain metabolites and responses to drug treatments (Hong et al. 2011, Neeley et al. 2011, Manduca et al. 2015). Furthermore, albino and pigmented rat strains display differences in baseline behaviours as well as drug responses, possibly linked to underlying differences in synaptic processing between albino and LH strains (Manahan-Vaughan 2000, McDermott and Kelly 2008, Ihalainen et al. 2016). These findings all suggest that differences among rat strains may provide valuable information regarding behavioural and neurobiological diversity relevant to variability in treatment responses among patients. Of course, if the underlying factors involved in the variation of responses to cannabinoid treatment are to be properly modelled in rodents, it would be crucial to also consider sex differences. Research in both humans and rodents suggest cannabinoids are more potent in females compared with males (Craft et al. 2013, Wei et al. 2017). In rodent models, male and female LH rats differ in brain CB1R density and function (Castelli et al. 2014) and in SD rats URB597 improves NOR only in adult male but not in adult female rats (Hlavacova et al. 2015). Taken together, these findings suggest that testing drugs in multiple rodent strains and in both females and males may reveal crucial information to understanding the variability in responses among patients.

## Conclusions

URB597 did not alter social preference and memory, NOR or the total sum of SI in LH rats, which is in contrast to findings in Wistar and SD rats, but in accord with observations of differences in responses to eCB modulation between albino and pigmented rat strains. Strain-dependent differences in laboratory rodents can be a source of frustration when designing and interpreting pre-clinical experiments. However, genetic variation between rodent strains may also provide opportunities to examine the behavioural and neurobiological differences in drug responses observed in the human population. In the present study, neither doses of URB597 affected the total sum of social interaction in LH rats, in contrast to findings in albino strains, but the higher dose of URB597 shifted the composition of social behaviour towards increased aggression. This finding underlines the importance of understanding the nature of drug-induced alterations in the expression of aggression.

## Abbreviations

2-AG: 2- arachidonoylglycerol
AEA: Anandamide
CB1R: Cannabinoid receptor type 1
CB2R: Cannabinoid receptor type 2
eCB: Endocannabinoid
FAAH: Fatty acid amide hydrolase
LH: Lister Hooded
NOR: Novel object recognition
SD: Sprague-Dawley
SI: Social interaction
SNT: Social novelty test
TRPV1: Transient receptor potential vanilloid 1

## Acknowledgements

WGW was supported by a Biotechnology and Biological Sciences Research Council (BBSRC) Doctoral Training Partnership [grant number BB/M008770/1] and the University of Nottingham. EPP was supported by a BBSRC Industrial CASE PhD studentship [grant number BB/M008770/1], which was co-sponsored by Artelo Biosciences. CS was supported by a research grant from the BBSRC [grant number BB/P001149/1]. The funders had no other role in any aspect of this paper.

## Contribution to the field

The endocannabinoid system has been implicated in both social and cognitive processing. The endocannabinoid metabolism inhibitor, URB597, dose-dependently improves non-social memory in albino rat strains, whereas its effect on social interaction (SI) is affected by both rat strain and drug dose. Lister Hooded rats consistently respond differently to drug treatment in general compared with albino strains. This study sought to investigate the effects of different doses of URB597 on social and non-social memory in Lister Hooded rats, as well as analysing the detailed behavioural composition of SI. Males were tested for novel object recognition (NOR), social preference, social novelty recognition and SI with an unfamiliar rat. Lister Hooded rats displayed expected non-social and social memory and social preference and SI, which was not affected by URB597. However, during SI, the high dose increased aggression, compared to vehicle, and decreased anogenital sniffing, compared to the low dose of URB597. These findings highlight the importance of considering strain and behavioural composition when investigating drug effects on social behaviour.

## Author Contributions

WGW and CS designed the study. WGW, EH, EPP and CS conducted the experiments. WGW, CWS and CS analysed the data and drafted the manuscript. WGW, HJC, CWS, and CS revised and approved the final version of the manuscript.

## Data Accessibility

In accordance with open science practices, all videos and analysis files are accessible at https://osf.io/zsm8f/

